# Diatoms structure the plankton community based on selective segregation in the world’s ocean

**DOI:** 10.1101/704353

**Authors:** Flora Vincent, Chris Bowler

**Affiliations:** lnstitut de Biologie de l’ENS (IBENS), Département de biologie, École normale supérieure, CNRS, INSERM, Université PSL, 75005 Paris, France

## Abstract

Diatoms are a major component of phytoplankton, believed to be responsible for around 20% of the annual primary production on Earth. As abundant and ubiquitous organisms, they are known to establish biotic interactions with many other members of the plankton. Through analysis of co-occurrence networks derived from the *Tara* Oceans expedition that take into account the importance of both biotic and abiotic factors in shaping the spatial distributions of species, we show that only 13% of diatom pairwise associations are driven by environmental conditions, whereas the vast majority are independent of abiotic factors. In contrast to most other plankton groups, at a global scale diatoms display a much higher proportion of negative correlations with other organisms, particularly towards potential predators and parasites, suggesting that their biogeography is constrained by top down pressure. Genus level analyses indicate that abundant diatoms are not necessarily the most connected, and that species-specific abundance distribution patterns lead to negative associations with other organisms. In order to move forward in the biological interpretation of co-occurrence networks, an open access extensive literature survey of diatom biotic interactions was compiled, of which 18.5% were recovered in the computed network. This result reveals the extent of what likely remains to be discovered in the field of planktonic biotic interactions, even for one of the best known organismal groups.

**Importance:** Diatoms are key phytoplankton in the modern ocean involved in numerous biotic interactions, ranging from symbiosis to predation and viral infection, which have considerable effects on global biogeochemical cycles. However, despite recent large-scale studies of plankton, we are still lacking a comprehensive picture of the diversity of diatom biotic interactions in the marine microbial community. Through the ecological interpretation of both inferred microbial association networks and available knowledge on diatom interactions compiled in an open access database, we propose an eco-systems level understanding of diatom interactions in the ocean.

## Introduction

Marine microbes, composed of bacteria, archaea and protists, play essential roles in the functioning and regulation of Earth’s biogeochemical cycles (Falkowski *et al.*, 2008). Their roles within planktonic ecosystems have typically been studied under the prism of bottom-up research, namely understanding how resources and abiotic factors affect their abundance, diversity and functions. On the other hand the effect of mortality, allelopathy, symbiosis and other biotic processes are also likely to shape their communities and to exert strong selective pressures on them (Strom, 2008; Smetacek, 2018), yet have been studied much less. With concentrations reaching 10^^7^/L protists (Brown *et al.*, 2009) and 10^^9^/L prokaryotes (Whitman *et al.*, 1998), biotic interactions are likely to impact community structure from the microscale to the ecosystem level (Zehr *et al.*, 2017)(Zehr and Weitz).

Among marine protists, diatoms (Bacillariophyta) are of key ecological importance. They are a ubiquitous and predominant component of phytoplankton, characterized by their ornate silica cell walls, and responsible for approximately 40% of marine net primary productivity (NPP; (Nelson *et al.*, 1995; Field, 1998)). The array of biotic interactions in which marine diatoms have been described is vast. They are fed upon by heterotrophic microzooplankton such as ciliates and phagotrophic dinoflagellates (Sherr and Sherr, 2007; Wang *et al.*, 2012; Zhang *et al.*, 2017), as well as by metazoan grazers such as copepods (Runge, 1988; Smetacek, 1998; Falkowski, 2002; Turner, 2014). Other known interactions include symbioses with nitrogen-fixing cyanobacteria (Foster and Zehr, 2006; Foster *et al.*, 2011) and tintinnids (Vincent *et al.*, 2018), parasitism by chytrids and diplonemids (Gsell *et al.*, 2013), diatom-targeted allelopathy by algicidal prokaryotes and dinoflagellates (Paul and Pohnert, 2011; Poulson-Ellestad *et al.*, 2014) and allelopathy mediated by diatom-derived compounds detrimental to copepod growth (Pohnert, 2005; Carotenuto *et al.*, 2014). Beyond direct biotic interactions, diatoms are also known to thrive in high nutrient and high turbulent environments such as upwelling regions, at the expense of the other major phytoplankton groups, for instance dinoflagellates and haptophytes (Margalef, 1979; Kemp and Villareal, 2018). Competition for silicon between diatoms and radiolarians, another silicifying member of the plankton, has also been noted (Harper and Knoll, 1975; Hendry *et al.*, 2018).

Despite the strong biotic and abiotic selective pressures that likely influence diatom biogeography and evolution, they are considered as successful r-selected species (Armbrust, 2009). r-selection is an evolutionary strategy in which species can quickly produce many offspring in unstable environments, at the expense of individual “parental investment” and low probability of surviving to adulthood. Rats are a classic example of r-selected species. This is opposed to K-selection, in which species produce fewer descendants with increased parental investment, such as elephants or whales (Pianka, 1970). Diatoms are one of the most diverse planktonic groups in terms of species, widely distributed across the world’s sunlit ocean (Malviya *et al.*, 2016) and capable of generating massive “blooms” in which diatom biomass can increase up to three orders of magnitude in just a few days (Platt *et al.*, 2009). Their success has been attributed, in part, to a broad range of predation avoidance mechanisms (Irigoien, 2005) such as their solid mineral skeleton (Hamm and Smetacek, 2007), chain and spine formation in some species, and toxic aldehyde production (Miralto *et al.*, 1999; Vardi *et al.*, 2006). However, a global view of their capacity to interact with other organisms and an assessment of its consequences on community composition are still lacking.

Co-occurrence networks using meta-omics data are increasingly being used to study microbial communities and interactions (Faust and Raes, 2012; Li *et al.*, 2016), e.g., in human and soil microbiomes (Barberán *et al.*, 2012; Faust *et al.*, 2012) as well as in marine and lake bacterioplankton (Fuhrman and Steele, 2008; Eiler *et al.*, 2012; Milici *et al.*, 2016). Such networks provide an opportunity to extend community analysis beyond alpha and beta diversity towards a simulated representation of the relational roles played by different organisms, many of whom are uncultured and uncharacterized (Proulx *et al.*, 2005; Chaffron *et al.*, 2010). Over large spatial scales, non-random patterns according to which organisms frequently or never occur in the same samples are the result of several processes such as biotic interactions, habitat filtering, historical effects, as well as neutral processes (Fuhrman, 2009). Quantifying the relative importance of each component is still in its infancy. However, these networks can be used to reveal niche spaces, to identify potential biotic interactions, and to guide more focused studies. Much like in protein-protein networks, interpreting microbial association networks also relies on literature-curated gold standard databases (Li *et al.*, 2016), although such references are woefully incomplete for most planktonic groups (Poelen *et al.*, 2014).

As part of the recent *Tara* Oceans expedition (Karsenti *et al.*, 2011; Bork *et al.*, 2015), determinants of community structure in global ocean plankton communities were assessed using co-occurrence networks (Lima-Mendez *et al.*, 2015), based on the abundance of viruses, bacteria, metazoans and algae across 68 *Tara* Oceans stations in two depth layers in the photic zone (**Annexe A**). Pairwise links between species were computed based on how frequently they were found to co-occur in similar samples (positive correlations) or, on the contrary, if the presence of one organism negatively correlated with the presence of another (negative correlations). In order to prevent spurious correlations due to the presence of additional confounding components such as abiotic factors, interaction information was furthermore calculated to assess whether or not edges were driven by an environmental parameter.

## Results

### Diatoms are segregators in the open ocean

Co-occurrence amongst different micro-organisms in the plankton has recently been investigated at a large scale using data from *Tara* Oceans by Lima-Mendez et al, 2015. The resulting interactome, denoted the *Tara* Oceans Interactome, represents species (nodes), connected by links (edges) that represent either positive or negative associations. Positive associations should be understood as two organisms that are often abundant in the same sample whereas negative associations emerge if the presence of one organism negatively correlates with the presence of another. Due to the potential impact of environmental drivers on the co-occurrence of two organisms, the *Tara* Oceans Interactome also provides insight into the effects of abiotic factors on pairwise correlations. This thus provides the opportunity to disentangle biotic from abiotic factors at the pairwise level.

The *Tara* Oceans Interactome has global coverage and reports over 90,000 statistically significant correlations, with ~68,000 of them being positive, ~26,000 of them being negative, and ~9,000 due to the simultaneous higher correlation of two organisms (OTUs) with a third environmental parameter. Diatoms are involved in 4,369 interactions, making them the 7th most connected taxonomic group after syndiniales (MALVs), arthropods, dinophyceae, polycystines, MASTs and prymnesiophyceae, independently from the taxon’s abundance (Lima-Mendez *et al.*, 2015). Overall, diatoms represent around 3% of all the positive associations (2,120/68,856) and 9.5% of all negative associations (2,249/23,777) showing that their contribution to negative associations is much higher than their contribution to positive co-occurrences, contrasting with all the other major taxonomic groups in the interactome. The positive to negative ratio of number of associations provides a measure of the group’s role in the network. Diatoms (Ratio = 0.99) and polycystines (Ratio = 0.66) are the only two groups that have more negative than positive associations and thus can be defined as “segregators” following the definition of Morueta-Holme, 2016. **(Table S1)**.

A finer analysis revealed the major taxonomic groups with which diatoms correlate or anti-correlate. Positive correlations involve mainly arthropoda (9.2% of diatom positive correlations), dinophyceae (8.7%), and syndiniales (an order of dinoflagellates also known as Marine Alveolates – “MALV” – found as parasites of crustacean, protists and fish) (11.7%). Negative correlations include the three previous groups – arthropoda (11.5%), dinophyceae (11.3%), syndiniales (11.1%) – as well as the polycystina (6%), a major group of radiolarians that produce mineral skeletons made from silica (**Figure 1.a**). Chlorophyceae were used as a control class for obligate photosynthetic green algae, and dictyochophyceae were used as a control class for silicified phytoplankton: both photosynthetic classes show more copresences with the aforementioned groups (**Figure 1.b-c**). However polycystines show similar exclusion trends as diatoms (**Figure 1.d**). The number of negative correlations involving diatoms with arthropods, dinophyceae, syndiniales and polycystines was much higher than what would be expected at random based on binomial testing (**Figure 1.e**), a pattern that was not found in other phytoplankton control groups. However, polycystines also display more negative associations than what would be expected at random with copepods, syndiniales and dinoflagellates **(Table S2)**. Amongst all the pairwise associations involving diatoms and other organisms in the plankton (N=4,369), only 13% were due to a third environmental parameter, illustrating a shared preference for a particular abiotic condition (N=566), leaving 87% of the associations solely explained by the abundance of the two organisms **(Figure 2 and Table S3)**. Polycystines displayed a similar pattern, with 95% of the association explained by biotic interactions rather than abiotic.

**Figure 1:**
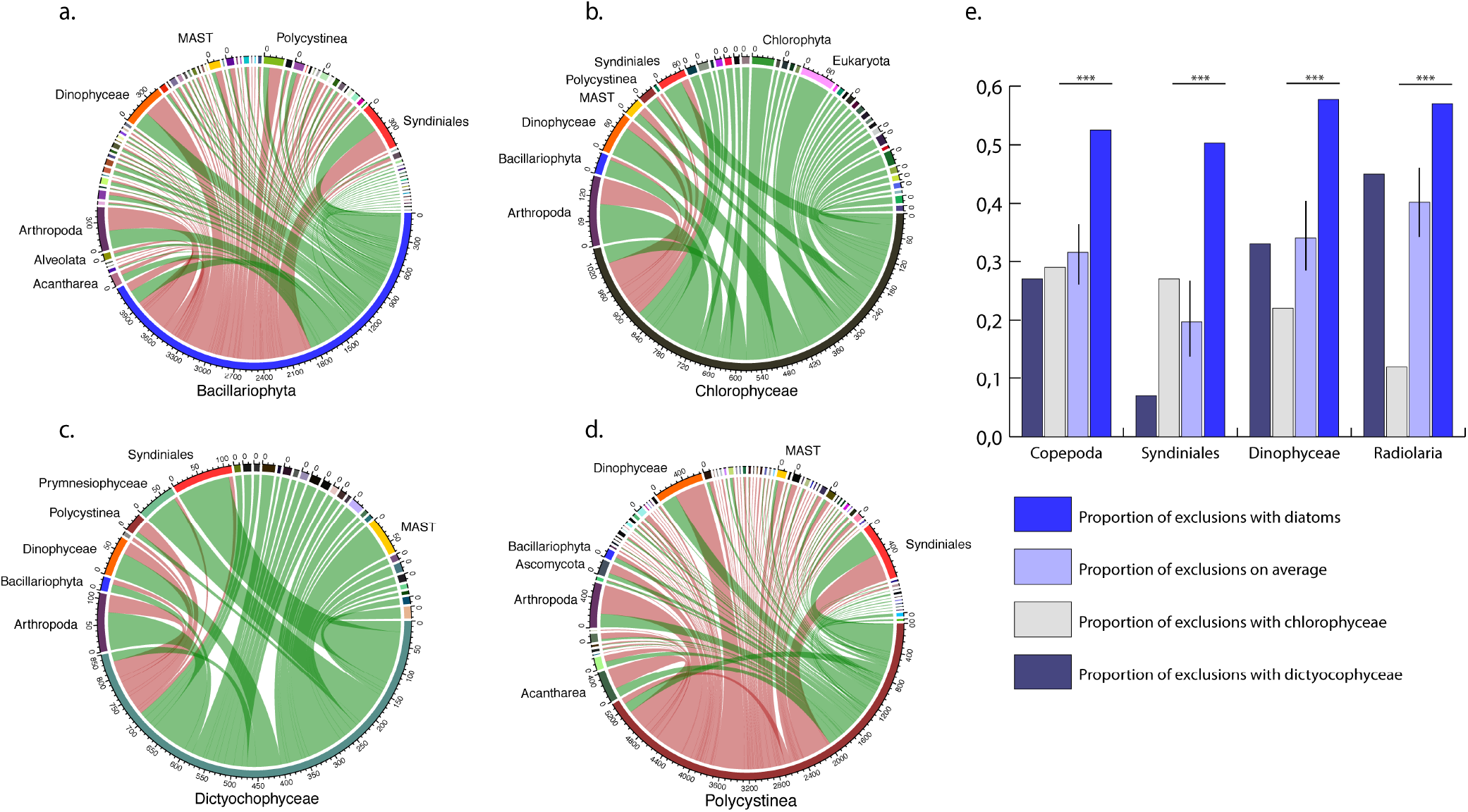
Major patterns of interactions for diatoms and control groups. Circos plots showing all copresences and exclusions of a) diatoms b) chlorophyceae (green algae control group), c) dictyocophyceae (biflagellates mixotroph silicifyer) and d) polycystines (the only other segregator). All size fractions networks are represented here. e) Comparison of proportion of exclusions showing diatoms significantly exclude potential predators, parasites and competitors such as Copepodes, Syndiniales, Dinophyceaes and Radiolarias, compared to control groups.

**Figure 2:**
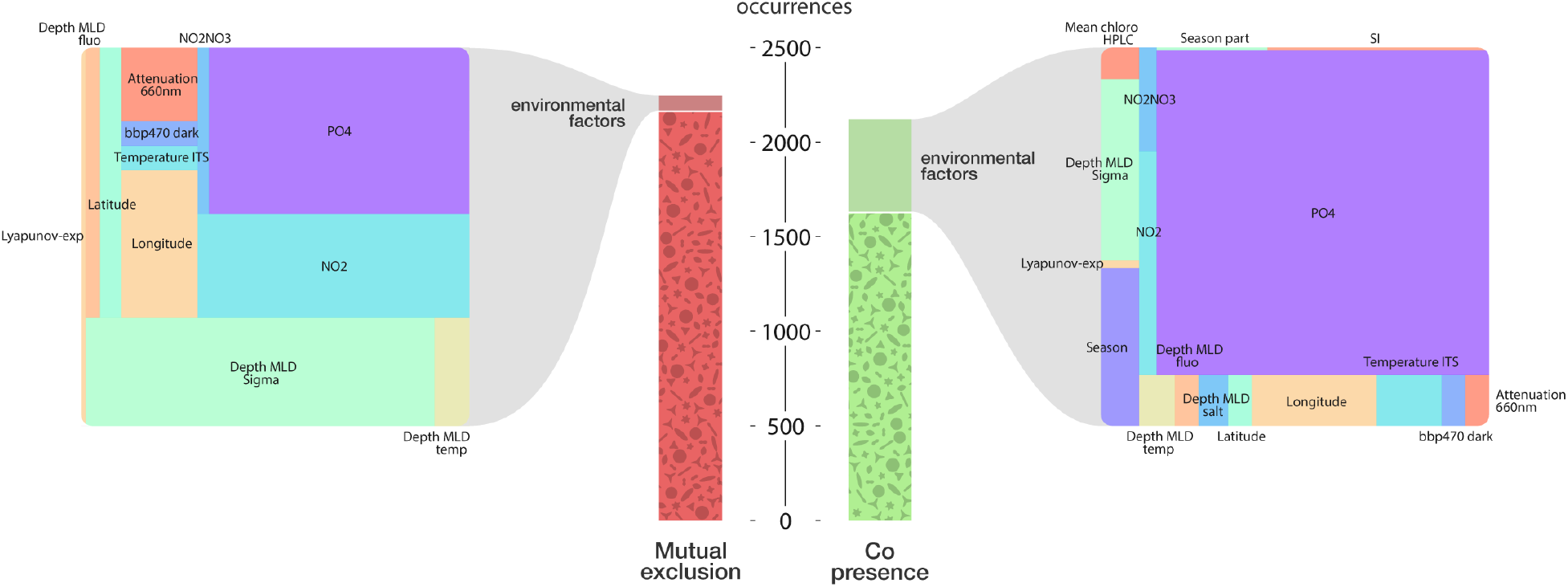
Biotic versus abiotic drivers of diatoms interactions. The methodology used to disentangle the relative influence of abiotic factors on diatom interactions is available in (Lima-Mendez *et al.*, 2015). Environmental Parameters: Phosphate (PO4); Mixed Layer Depth (MLD, layer in which active turbulence homogenizes water, estimated by density – sigma- and temperature); Nitrite (NO2); Light scattering by suspended particles (Beam attenuation 660nm); Backscattering coefficient of particle (bbp470); HPLC Chlorophyll pigment measurement (HPLC-adjusted); Ocean perturbation (Lyapunov exponent), Silicate (SI) and categorical variable for season. A full description of the environmental parameters is available on the PANGAEA website (https://doi.pangaea.de/10.1594/PANGAEA.840718).

Sub-networks were then extracted for both positive and negative associations involving diatoms with copepods, dinophyceae, syndiniales (MALVs) and MASTs (a group of small, flagellated, bacterivorous stramenopiles) **(Figure 3.a)**. The size of each node corresponds to a continuous mapping of the betweenness centrality, representative of the importance of a node in maintaining the overall structure of a network. What is noteworthy is that important diatom genera are not always the same depending on the partner of interaction. More specifically, Syndiniales and Dinophyceae sub-networks involve mainly *Pseudo-nitzschia*, *Actinocyclus*, and *Chaetoceros* assigned barcodes, whereas important nodes in copepods and MASTs involve *Thalassosira, Leptocylindrus or Synedra*, showing a nonrandom pattern of species co-occurrence. Many MAST nodes belong to the MAST-3 clade, known to harbor the diatom parasite *Solenicola setigara* (Gómez, 2011). In order to compare the architecture between the four sub-networks, we investigated specificity of the interaction, asking if all organisms are interconnected using topological metrics such as connected components. Diatom-MALV and diatom-MAST sub-networks have more connected components suggesting more specialist interactions than diatoms with copepods or dinophyceaes **(Figure 3.b and Table S4)** and average scores of exclusions were stronger for diatom-MAST (−0.66+−0.09) and diatom-MALV (−0.59+−0.09) subnetworks (**Figure 3.c**).

**Figure 3:**
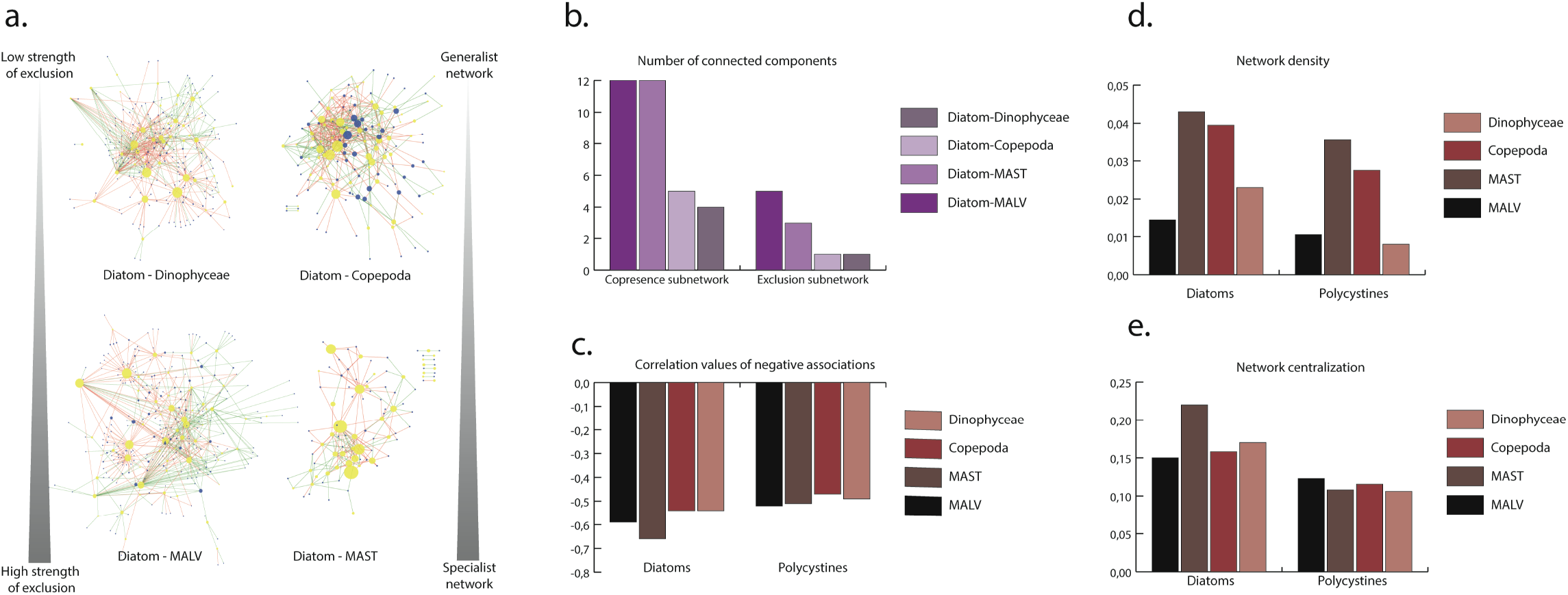
Subnetworks of diatom with major interacting groups. a) Diatom nodes are colored in yellow, and the corresponding partner nodes are colored in blue. Green edges correspond to positive co-occurrences while red edges correspond to negative correlations. The size of the node corresponds to a continuous mapping of the degree in the global diatom interactome. b) Number of connected components within each diatom network, separated by copresence or exclusion. Comparison of c) exclusion correlation values, d) network density and e) network centralization between diatoms and polycystines with their major partners of interaction.

We used polycystines as a comparison group as they were also shown to be segregators. Diatoms have stronger negative scores than polycystines, reflecting a higher potential as segregators with respect to potential competitors, grazers and parasites such as copepods, dinophyceae, and syndiniales **(Figure 3.c)**. Furthermore, diatoms tend to form much denser (more interconnected i.e., less species-specific (**Figure 3.d**) and centralized (relying on fewer central species (**Figure 3.e**)) networks than polycystines. Despite comparable patterns of segregation between diatoms and polycystines, they differ in strength of negative interactions based on Spearman correlation values and how specific the interactions are at the barcode level.

### Global-scale genus abundance does not determine importance in connectivity

While abundant diatoms are likely to be important players in biogeochemical cycles such as NPP and carbon export, how their biotic interactions influence plankton community diversity and abundance is still unknown. To address this question, the ten most abundant diatom genera, defined based on 18S V9 read abundances (Malviya *et al.*, 2016), were analyzed with respect to their positions in the diatom interactome **(Table S5)**. This analysis revealed that some barely play a role in the interactome. For example, *Chaetoceros* is the most abundant genus (1,615,027 reads), yet it is only represented in 515 edges across the interactome. Hence, no significant correlation was found between the total abundance of the genus and the number of edges (i.e., putative biotic relations the genus is involved in) (Spearman p.value = 0.96), nor the number of nodes involved (i.e., the number of different interacting organisms) (Spearman p.value = 0.45) (**Figure S1**). On the other hand, the diatom genus *Synedra*, that is not abundant at the global level (ranked as the 22^nd^ most abundant diatom with 28,700 reads), was involved in over 100 significant associations. *Pseudo-nitzschia* is the top assigned co-occurring diatom representing 7% of the positive interactions in the diatom network; on the contrary, exclusions involved a large array of diatom genera each representing on average 2% of the interactions, suggesting exclusion to be a shared property amongst diatoms **(Figure S2)**.

Statistics of network level properties provide further insights into the overall structure of genera-specific assemblages, and was investigated at the genus level for the most connected ones **(Figure S3 and Table S6)**. *Leptocylindrus*, *Proboscia*, and *Pseudo-nitzschia* displayed a higher average number of neighbors, meaning their sub-networks are highly interconnected between diatom and non-diatom OTUs, suggesting that interactions within those genera are not species specific. On the other hand, the *Chaetoceros*, *Eucampia*, and *Thalassiosira* sub-networks displayed larger diameters, meaning that a few diatom OTUs are connected both positively and negatively to a large number of partners that are not connected to any other diatom OTUs, indicative of a more species-specific type of behavior with respect to interactions. No clear correlation was found between the crown age estimation of marine planktonic diatoms nor taxon richness estimated from the number of OTU swarms (Nakov *et al.*, 2018), and the number of associations they are involved in **(Table S6)**, suggesting that the establishment of biotic interactions is a continuous and dynamic process independent of the age of a diatom genus.

### Species level segregation determined by endemic and blooming diatoms

Due to the small number of individual barcodes in the interactome that have species level resolution, we decided to conduct a finer analysis and ask whether or not different barcodes of the same (abundant) genera display specificity in the type of interactions and partners they interact with. We illustrate this barcode specificity with three different examples: *Chaetoceros*, *Pseudo-nitzschia* and *Thalassiosira*. *Chaetoceros* interactions reveal that different species display very different co-occurrence patterns. The barcode “29f84,” assigned to *Chaetoceros rostratus*, is essentially involved in positive co-occurrences, while barcode “8fd6d” assigned to *Chaetoceros debilis*, is the major driver of negative associations involving dinophyceae, MASTs, syndiniales and arthropods **(Figure 4.a)**. This could reflect the different species tolerance to other organisms, since several *Chaetoceros* species are known to be harmful to aquaculture industries (Albright *et al.*, 1993); *Chaetoceros debilis* in particular can cause physical damage to fish gills (Kraberg *et al.*, 2010).

**Figure 4:**
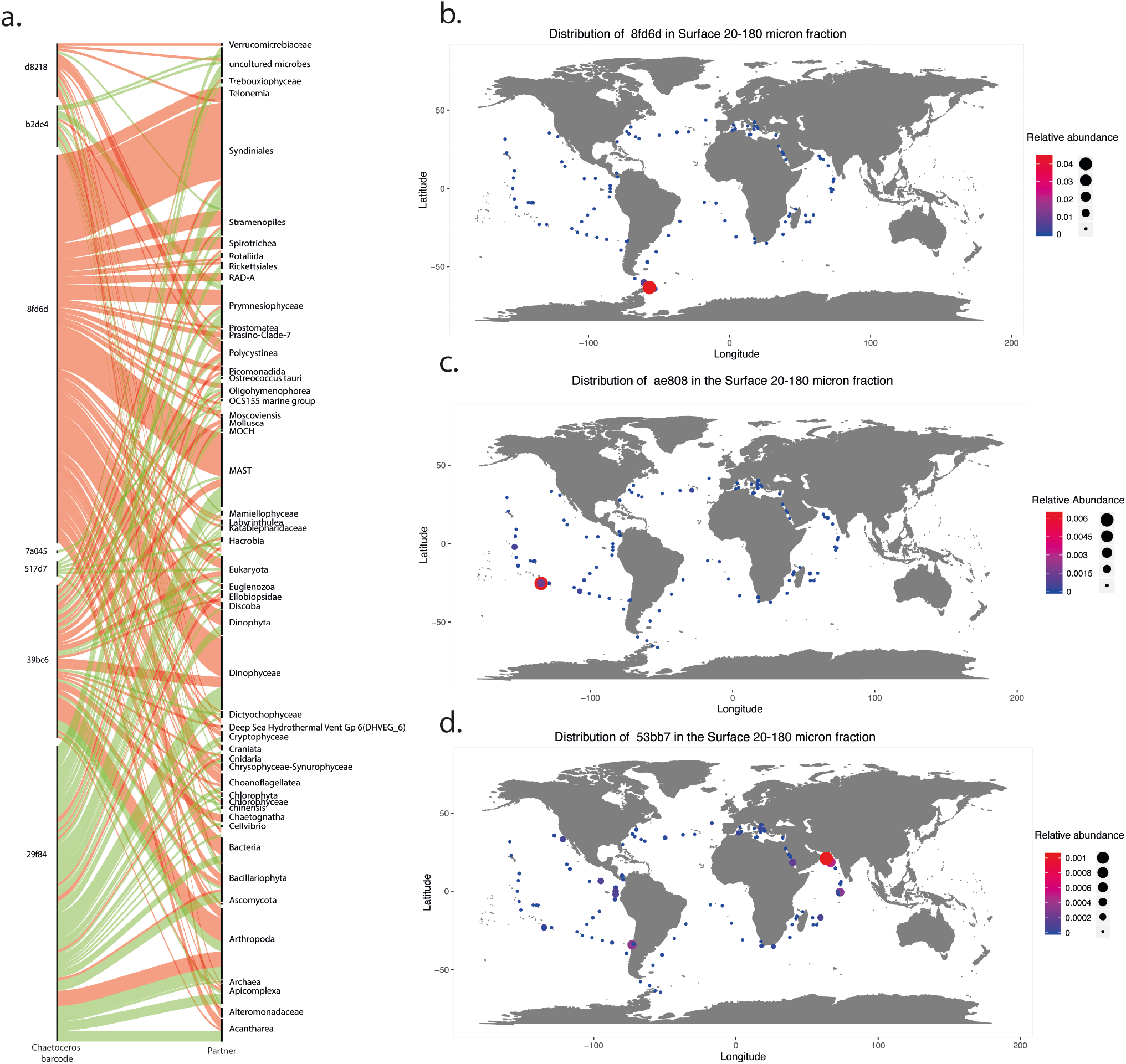
Barcode level specificity of interactions and biogeographic distribution of top excluding diatoms. a) Barcode level associations of the diatom genus *Chaetoceros*. Diatom barcodes annotations are listed on the left, and partners of interaction on the right. The thickness of the band corresponds to the number of interactions, with copresences in green and exclusions in red. 29f84, 7a045: *Chaetoceros rostratus*; 8fd6d, b2de4 *Chaetoceros sp*.; 39bc6, 517d7, d8218: *Chaetoceros muellerii*. Biogeography of top excluding diatom barcodes in surface waters of the 20-180 micron fraction b) 8fd6d: *Chaetoceros rostratus* c) ae808: *Thalassiosira sp*. d) 53bb7 : *Proboscia sp.*.

*Pseudo-nitzschia* barcodes are primarily involved in positive correlations. However, they display exclusions with organisms such as arthropoda and dinophyceae, and some are known to produce the toxin domoic acid in specific conditions (Tammilehto *et al.*, 2015). No exclusions regarding syndiniales appear, and barcode-level specificity is observed with “1d16c,” which is involved in a much higher number of interactions than “b56c3.” Unfortunately, these diatom sequences were not assigned at the species level. Finally, the *Thalassiosira* sub-network displays mostly negative associations with syndiniales, arthropoda, and polycystines, with one of the three representative barcodes (“53bb7”) responsible for 93% of the exclusion (**Table S6**). The distribution of the aforementioned diatoms, involved in a high number of mutual exclusions, is typical to that of endemic and blooming diatoms as their read abundance massively increases either in specific localized stations, or nutrient replete well mixed regions. This observation was supported by analyzing the distribution patterns of the 6 top diatom barcodes involved in exclusions (**Table S6**) such as 90dad (226 exclusions – Unassigned *Bacillariophyta* blooming in Indian Ocean Station TARA_036), 4c4a8 (193 exclusions – *Raphid-Pennate*, Marquesas station TARA_122), 8fd6d (168 exclusions – *Chaetoceros* in Southern Ocean Station TARA_088, **Figure 4.b**), 30191 (166 exclusions – *Actinocyclus* in Indian Ocean Station TARA_033), 53bb72 (94 exclusions – *Thalassiosira* in Indian Ocean Station TARA_036 **Figure 4.c**), ae808 (103 exclusions – *Proboscia*, Station TARA_116, **Figure 4.d**).

### Diatom – bacteria interactions in the open ocean

Diatom-prokaryote associations represent 830 interactions, or 19% of the whole diatom co-occurrence network (**Table S7**). This can be considered as average when compared to bacteria associations in copepod interactions (28%), dinophyceae (18.5%), radiolaria (20.5%) and syndiniales (16.3%). By classifying the bacteria according to their primary nutritional group (see Methods), diatoms were found to be more associated, both positively and negatively, to heterotrophs (637 associations) than to autotrophs (87 associations) (**Figure S4**). Even though diatoms do not significantly co-occur with or exclude a specific bacterial nutritional group, many exclusions involve Rhodobacteraceae, SAR11 and SAR86 clades **(Figure 5)**. Interestingly, diatom specific patterns are apparent. For example, the *Actinocyclus* and *Haslea* diatom genera are solely involved in exclusions against a wide range of bacteria, whereas *Pseudo-nitzschia* is mainly involved in co-presences. Interestingly, *Haslea ostrearia* is known for producing a water-soluble blue pigment, marennine, of which closely related pigments display antibacterial activities (Gastineau *et al.*, 2014).

**Figure 5:**
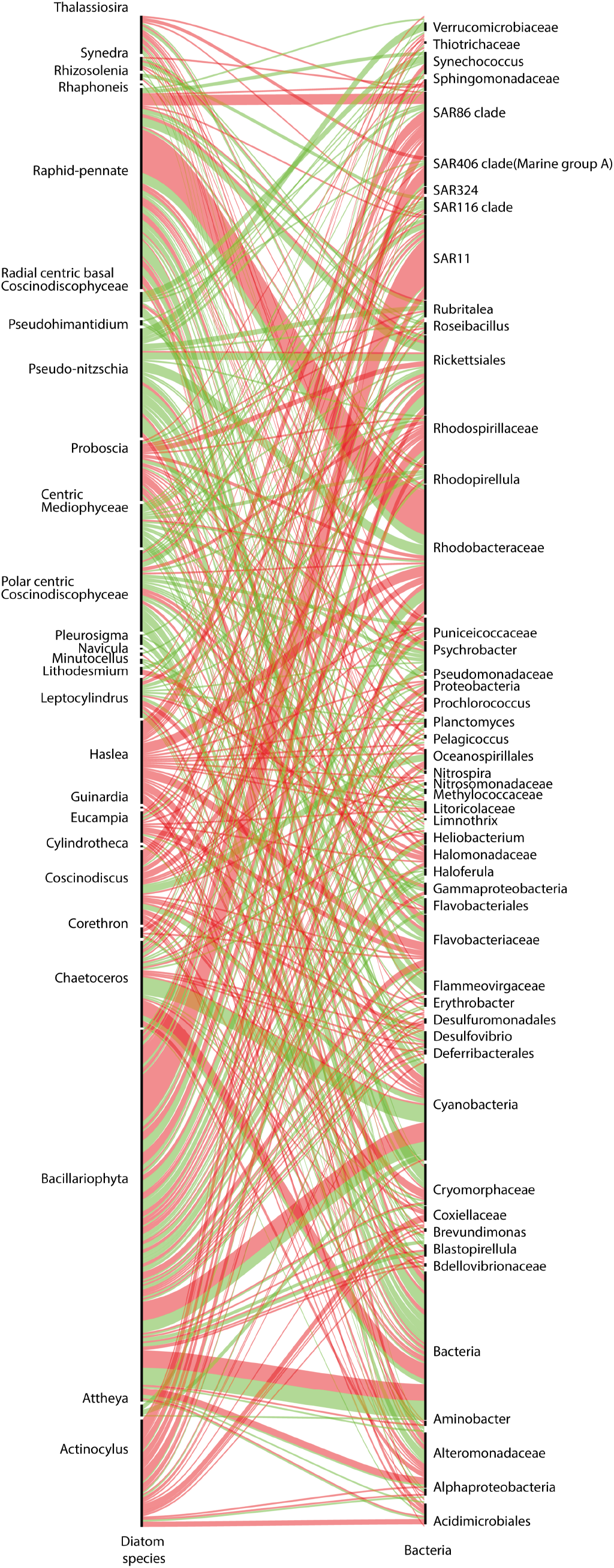
Diatom bacteria interactions. Diatom taxonomic annotations are listed on the left, and partners of interaction on the right. The thickness of the band corresponds to the number of interactions, with copresences in green and exclusions in red.

### A skewed knowledge about diatom biotic associations

To review current knowledge about diatom interactions we generated an online open access database (DOI:10.5281/zenodo.2619533) that assembled the queryable knowledge in the literature about diatom associations from both marine and freshwater habitats, and is synchronized with GloBI, a global effort to map biological interactions (Poelen *et al.*, 2014). A total of 1,533 associations from over 500 analyzed papers involving 83 unique genera of diatoms and 588 unique genera of other partners are reported here and made available through the GloBi database, illustrating a diversity of association types such as predation, symbiosis, allelopathy, parasitism, and epibiosis, as well as the diversity of partners involved in the associations, including both prokaryotes and eukaryotes, micro- and macro-organisms **(Figure 6.a)**.

**Figure 6:**
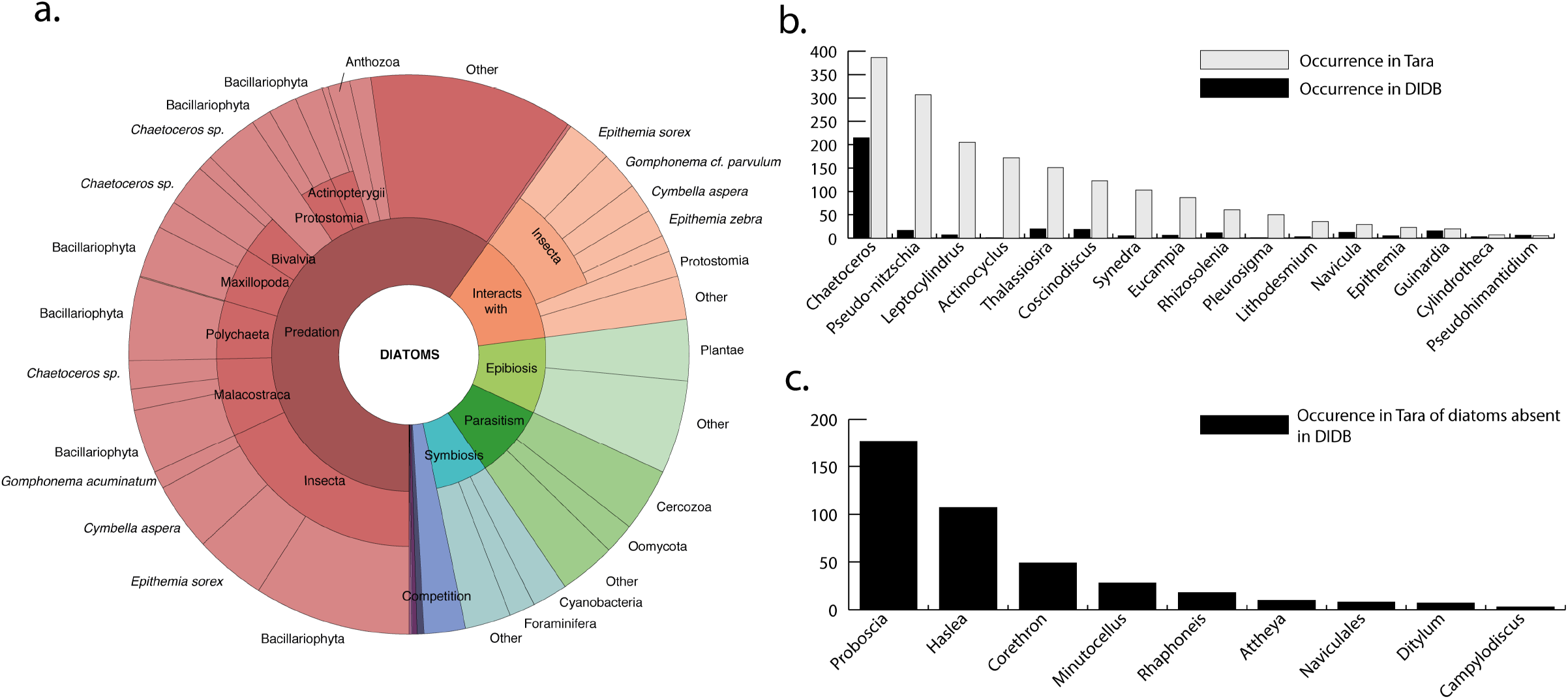
Current knowledge of diatom biotic interactions and comparison with the*Tara* Oceans Interactome. a) KRONA plot based on available literature concerning diatom associations, mined and manually curated from Web Of Science, PubMed and Globi. The outer circle represents the diatom genera (when known), the middle circle represent the interacting partner, and the inner circle represents the type of interaction (predation, parasitism, symbiosis) b) Comparison between number of interactions involving a specific diatom genus in the literature (black) and in the *Tara* Oceans Interactome (grey) showing strong disparities for diatoms such as *Pseudo-nitzschia* c) Number of interactions of important diatom genera in the interactome that are absent from the literature, suggesting interesting areas for future research.

We noted that 58% (883 out of 1,533) of the interactions are labelled “eatenBy” (“Predation” in **Figure 6.a**) and involve mainly insects (267 interactions; 30% of diatom predators) and crustaceans (15% of diatom predators). Cases of epibiosis, representing approximately 10% of the literature database, were largely dominated by epiphytic diatoms living on plants (40% of epibionts) and epizoic diatoms living on copepods (9% of epibionts). Parasitic and photosymbiotic interactions, although known to have significant ecological implications at the individual host level as well as at the community composition scale (Van Veen *et al.*, 2008), represented only 15% of the literature database for a total of 219 interactions, involving principally diatom associations with radiolarians and cyanobacteria. Interactions involving bacteria represent 72 associations (4.8 % of the literature database).

The distribution of habitats amongst the studied diatoms reveals a singular pattern: the majority of diatom interactions in the literature are represented by a handful of freshwater diatoms, whereas many marine species are reported in just a small number of interactions **(Figure S5)**. In terms of partners involved (detailed in **Figure S6)**, one third are represented by insects feeding upon diatoms in streams, and crustaceans feeding upon diatoms in both marine and freshwater environments. Other principal partners are plants, upon which diatoms attach as “epiphytes,” such as Posidonia (seagrass), Potamogeton (pondweed), Ruppia (ditchgrass) and Thalassia (seagrass). Consequently, our knowledge based on the literature produces a highly centralized network containing a few diatoms mainly subject to grazing or epiphytic on macro-organisms. Major diatom genera for which interactions are reported in the literature are *Chaetoceros spp* (215 interactions, marine and freshwater), *Epithemia sorex* (135 interactions, freshwater), and *Cymbella aspera* (115 interactions, freshwater).

### Overlapping empirical evidence from data-driven results reveals gaps in knowledge and extends it to the global ocean

In an effort to improve edge annotation in the co-occurrence network, the literature database presented here was used. The occurrence of a specific genera in the literature was compared to its occurrence in the *Tara* Oceans Interactome (**Table S8**). On average, the co-occurrence network revealed many more potential links between species than has been reported in the literature (**Figure 6.b**). Disparity was especially high for *Pseudo-Nitzschia*, mentioned in 17 interactions in the literature compared to 307 associations in the interactome. On the other hand, many diatoms involved in several associations in the interactome are absent from the literature, such as *Proboscia* and *Haslea* (**Figure 6.c**).

Of 1,533 literature-based interactions, 178 could potentially be found in the *Tara* Oceans Interactome, as both partners had a representative barcode in the *Tara* Oceans database. A total of 33 literature-based interactions (18.5% of the literature associations) were recovered in the network at the genus level, representing a total of 289 interactions from the interactome and 209 different barcodes. These 289 interactions represent 6.5% of all the associations involving Bacillariophyta in the *Tara* Oceans co-occurrence network. By mapping available literature on the co-occurrence network, we can see that the major interactions recovered are those involving competition, predation and symbiosis with arthropods, dinoflagellates and bacteria. However, predation by polychaetes and parasitism by cercozoa and chytrids are missing from the *Tara* Oceans interactome.

## Discussion

The *Tara* Oceans Interactome represents an ideal case study to investigate global scale community structure involving diatoms, as it maximizes spatio-temporal variance across a global sampling campaign and captures systems-level properties. Here, we reveal that diatoms and polycystines are the organismal groups with the highest proportion of negative associations within the *Tara* Oceans Interactome, and classify them as segregators according to a definition from (Morueta-Holme *et al.*, 2016), as they display more negative than positive associations. Diatoms and polycystines prevent their co-occurrence with a range of potentially harmful organisms over broad spatial scales (Smetacek, 2012) (**Figure 1.a,d**), a pattern unseen in the other photosynthetic classes examined (**Figure 1.b,c**) reflected by significant exclusion of major functional groups of predators, parasites and competitors such as Copepods, Syndiniales and Dinophyceae (**Figure 1.e**). Diatoms are known to have developed an effective arsenal composed of silicified cell walls, spines, toxic oxylipins, and chain formation to increase size, so we propose that the observed exclusion pattern reflects the worldwide impact of the diatom arms race against potential competitors, grazers and parasites. Additionally, building upon the phylogenetic affiliation of individual sequences, barcodes can be assigned to a plankton functional type that refers to traits such as the trophic strategy and role in biogeochemical cycles (Le Quéré *et al.*, 2005). As demonstrated in the *Tara* Oceans Interactome (Lima-Mendez *et al.*, 2015), diatoms compose the “phytoplankton silicifiers” metanode, and display a variety of mutual exclusions that again distinguish them from other phytoplankton groups. The role of biotic interactions is emphasized by the fact that out of the complete diatom association network, co-localization and co-exclusion of diatoms with other organisms are due to shared preferences for an environmental niche in 13% of the cases, emphasizing the importance of biotic factors in 87% of the associations (**Figure 2**).

Connectivity values of sub-network topologies suggest that diatom-MAST and diatom-MALV networks display more specialist interactions than diatom-copepod and diatom-dinophyceae networks (**Figure 3.b**). Correlation values are often neglected in co-occurrence analysis but here they reveal stronger exclusion patterns of diatoms against MASTs and MALVs (**Figure 3.c**). Exclusion between diatoms and MASTs is therefore more specific, and stronger, than compared to copepods or dinoflagellates. These properties are conserved in the other segregator group, polycystines. Yet diatoms outcompete polycystines with higher strengths of exclusions based on correlation values and denser networks suggesting more species-specific interactions in Polycystines (**Figure 3.c-e**). Previous work exploring abundance patterns amongst planktonic silicifiers in the *Tara* Oceans data (Hendry *et al.*, 2018) revealed strong size-fractionated communities: if the smallest size fraction (0.8-5 micron) contained a large diversity of silicifying organisms in nearly constant proportions, co-occurrence of diatom and polycystines was rare in bigger size fractions (20-180 micron), where the presence of one organism appeared to exclude the other.

Analysis at the genus level shows that abundant diatoms such as *Attheya* do not play a central role in structuring the community, contrary to *Synedra* that, at a global scale, is less significant in terms of abundance but is highly connected to the plankton community. We show the existence of a species level segregation effect that can be attributed to harmful traits (Kraberg *et al.*, 2010) (**Figure 4.a**), reflected by blooming and endemic distribution patterns for the top segregating diatoms (**Figure 4.b-d**). Although diatom blooms can sometimes be triggered by light and nutrient perturbation, very few of the negative associations were found to be driven by the measured environmental parameters, emphasizing the importance of the biotic component for the segregative effect. These results support previous observations indicating the importance of biotic interactions in governing ocean planktonic blooms and distribution (Irigoien, 2005; Behrenfeld and Boss, 2014).

Our literature survey reveals a skewed knowledge, focusing on freshwater diatoms and interactions with macro-organisms, with very few parasitic, photosymbiotic or bacterial associations (**Figure 6.a**). The relative paucity of marine microbial studies can be explained by the difficulty of accessing these interactions in the field, which obviously limits our understanding of how such interactions structure the community at global scale. Comparing empirical knowledge and data-driven association networks reveal understudied genera such as *Leptocylindrus* and *Actinocyclus*, and those that are not even present in the literature, such as *Proboscia* and *Haslea* (**Figure 6.b,c**). However, *Proboscia* is a homotypic synonym of *Rhizosolenia* that is found in the interactome, which illustrates the consequences of non-universal taxonomic denominations on diversity analysis.

While 18.5% of the literature database was recovered in the interactome, it explained only 6.5% of the 4,369 edges composing the diatom network. The gap between the 20% of diatom-bacteria interactions in the *Tara* Oceans Interactome compared to only 4.8% for diatom-bacteria associations described in the literature highlights how little we know about host-associated microbiomes at this stage. Most of the experimental studies focus on symbiosis with diazotrophs and roseobacters, and antibacterial activity of *Skeletonema* against bacterial pathogens. In many ways, this high proportion of unmatched interactions should be regarded as the “Unknown” proportion of microbial diversity emerging from metabarcoding surveys. Part of it is truly unknown and new, part of it is due to biases in data gathering and processing, and part of it is due to the lack of an extensive reference database.

Many challenges remain regarding the computation, analysis, and interpretation of co-occurrence networks despite their potential to uncover processes governing diatom-related microbial communities. Recent studies are exploring the methodological bias of each co-occurrence method by attempting to detect specific associations within mock communities (Weiss *et al.*, 2016) whilst others admit that applying network statistics to microbial relationships is subject to high variability depending on taxonomic level of the study and criteria used to compute networks (Williams *et al.*, 2014). Assigning biological interactions such as predation, parasitism or symbiosis to correlations is still cumbersome and will require both proper references of biotic interactions (Li *et al.*, 2016), and further studies that investigate dynamics of interactions through space and time, and their sensitivity to co-occurrence detection. Attempts to develop a theoretical framework for the detection of species interactions based on co-occurrence networks is underway (Morales-Castilla *et al.*, 2015; Cazelles *et al.*, 2016; Morueta-Holme *et al.*, 2016), but must be confronted with actual *in situ* data. Furthermore, a vast body of literature already exists in the field of ecological networks, traditionally focusing on observational non-inferred data and the modeling of foodwebs, host-parasite and plant-pollinator networks (Ings *et al.*, 2009; Bascompte, 2010). Various properties linked to the architecture of these antagonistic and mutualistic networks have been formalized, such as nestedness, modularity, or the impact of combining several types of interactions in a single framework (Thebault and Fontaine, 2010; Fontaine *et al.*, 2011). Enhanced cross-fertilization between the disciplines of ecological networks and co-occurrence networks would highly benefit both communities, ultimately helping to understand the laws governing the “tangled bank” (Darwin, 1859).

Diatoms have undoubtedly succeeded in adapting to the ocean’s fluctuating environment, shown by recurrent, predictable and highly diverse bloom episodes (Guillard and Kilham, 1977). They are considered as r-selected species with high growth rates under favorable conditions that range from nutrient-rich highly turbulent environments to stratified oligotrophic waters (Margalef, 1978; Alexander *et al.*, 2015; Kemp and Villareal, 2018). Their success has long been attributed to this ecological strategy; here we suggest that abiotic factors alone are not sufficient to explain their ecological success. The current study shows that diatoms have evolved to avoid co-occurring with potentially harmful organisms such as predators, parasites and pathogens (Smetacek, 2012), shedding light on the top down forces that could drive diatom evolution and adaptation in the modern ocean.

## Material and methods

### Relative proportion of co-occurrences and exclusions with respect to major partners and network analysis

All analyses were performed on the published co-occurrence network in Lima-Mendez, 2015. Environmental drivers of diatom related edges are available in **Table S3**. Four independent matrices were created from the interactome regarding the major partners interacting with diatoms (copepods, dinophyceae, syndiniales and radiolaria) containing only pairwise interactions that involved the major partner and binomial testing was done using the dbinom and pbinom function as implemented in the {stats} package of R version 3.3.0. Subnetwork topologies were analyzed using the NetworkAnalyzer plugin in Cytoscape (Shannon *et al.*, 2003) as described in (Doncheva *et al.*, 2012). Network topologies for major groups are available in **Table S4**.

### Major diatom interactions

The 10 most abundant diatom genera in the surface ocean were selected based on the work published by (Malviya *et al.*, 2016). Their co-occurrence network was extracted **(Table S5)** from the global interactome and analyzed at the ribotype level. Network topologies are available in **Table S6**. Distribution of individual barcodes was assessed across the 126 *Tara* Oceans sampling stations.

### Construction of Diatom Interaction Literature Data Base

Literature was screened until November 2017 to look for all ecological interactions involving diatoms to establish the current state of knowledge regarding the diatom interactome, both in marine and freshwater environments and is made available at the DOI: 10.5281/zenodo.2619533. It is designed to be completed by external contributions. Diatom ecological interactions as defined in this paper are a very large group of associations, characterized by (i) the nature of the association defined by the ecological interaction or the mechanism (predation, symbiosis, mutualism, competition, epibiosis), (ii) the diatom involved, and (iii) the partners of the interaction.

The protocol to build the list of literature-based interactions was the following (i) collect publications involving diatom associations using (a) the Web of Science query TITLE: (diatom*) AND TOPIC : (symbio* OR competition OR parasit* OR predat* OR epiphyte OR allelopathy OR epibiont OR mutualism); (b) Eutils tools to mine Pubmed and extract ID of all publications with the search url http://eutils.ncbi.nlm.nih.gov/entrez/eutils/esearch.fcgi?db=pubmed&term=diatom+symbiosis&usehistory=y and the same keywords; (c) the get_interactions_by_taxa(sourcetaxon = “Bacillariophyta”) function from the RGlobi package (Poelen *et al.*, 2014), the most recent and extensive automated database of biotic interactions; and (d) personal mining from other publication browsers and input from experts (ii) extract when relevant the partners of the interactions based on the title and on the abstract for Web of Science, Pubmed and personal references and normalize the label of the interaction based on Globi nomenclature (iii) display KRONA plot with Type of Interaction / Partner Class / Diatom genus / Partner genus_species **(Figure 6a)**. Cases of episammic (sand) and epipelon (mud) interactions were not considered as they involved association with non-living surfaces.

### Comparison of literature interactions and diatom interactome

All partner genera interacting with diatoms based on the literature were searched for in the *Tara* Oceans dataset based on the lineage of the barcode. For each barcode that had a match, identifiers (“md5sum”) were extracted creating a list of 954110 barcodes to be searched for in the global interactome (**Table S8**).

## Aknowledgments

CB acknowledges funding from the French Government “Investissements d’Avenir” programs OCEANOMICS (ANR-11-BTBR-0008), MEMO LIFE (ANR-10-LABX-54), and Paris Sciences et Lettres Research University (ANR-11-IDEX-0001-02), as well as the European Union Framework Programme 7 (MicroB3/No.287589), European Research Council Advanced Awards Diatomite and Diatomic under the European Union’s Horizon 2020 research and innovation programme (grant agreement Nos. 294823 and 835067), the LouisD Foundation of the Institut de France, and a Fellowship from the Radcliffe Institute for Advanced Study at Harvard University. FJV acknowledges the Fondation de la Mer. This article is contribution # of *Tara* Oceans.

